# Proteomics reveals commitment to germination in barley seeds is marked by loss of stress response proteins and mobilisation of nutrient reservoirs

**DOI:** 10.1101/2020.06.02.130823

**Authors:** Sarah K. Osama, Edward D. Kerr, Adel M. Yousif, Toan K. Phung, Alison M. Kelly, Glen P. Fox, Benjamin L. Schulz

## Abstract

Germination is a critical process in the reproduction and propagation of flowering plants, and is also the key stage of industrial grain malting. Germination commences when seeds are steeped in water, followed by degradation of the endosperm cell walls, enzymatic digestion of starch and proteins to provide nutrients for the growing plant, and emergence of the radicle from the seed. Dormancy is a state where seeds fail to germinate upon steeping, but which prevents inappropriate premature germination of the seeds before harvest from the field. This can result in inefficiencies in industrial malting. We used Sequential Window Acquisition of all THeoretical ions Mass Spectrometry (SWATH-MS) proteomics to measure changes in the barley seed proteome throughout germination. We found a large number of proteins involved in desiccation tolerance and germination inhibition rapidly decreased in abundance after imbibition. This was followed by a decrease in proteins involved in lipid, protein and nutrient reservoir storage, consistent with induction and activation of systems for nutrient mobilisation to provide nutrients to the growing embryo. Dormant seeds that failed to germinate showed substantial biochemical activity distinct from that of seeds undergoing germination, with differences in sulfur metabolic enzymes, endogenous alpha-amylase/trypsin inhibitors, and histone proteins. We verified our findings with analysis of germinating barley seeds from two commercial malting facilities, demonstrating that key features of the dynamic proteome of germinating barley seeds were conserved between laboratory and industrial scales. The results provide a more detailed understanding of the changes in the barley proteome during germination and give possible target proteins for testing or to inform selective breeding to enhance germination or control dormancy.

## Introduction

Barley (*Hordeum vulgare* L. subsp. *vulgare*) is one of the most important cereal crops, ranked fourth highest in terms of production globally and is used in the production of beer, as livestock feed, and for human consumption [1]. For beer brewing, barley must first be malted, which consists of three stages: steeping, germinating, and kilning [2,3]. The key to malting is germination, in which the radicle emerges from the seed following imbibition of water [4]. As the radicle begins to emerge, the endosperm cell wall is degraded, providing newly synthesised enzymes access to the biopolymers within, enabling degradation of starch and proteins [4–6]. Barley starch consists of two polysaccharides, long chains of amylose and branched chains of amylopectin [7]. α- and β-amylase, along with other enzymes, digest these starch molecules into smaller useable sugars [7,8]. In addition, proteases break down seed storage and other proteins into small peptides and amino acids, which along with the small sugars can be used as nutrients by the developing embryo [9].

Prior to germination, barley seeds proceed through seed development, grain filling, and maturation. During grain filling and maturation there is an accumulation of storage proteins, lipids, and polysaccharides in the endosperm [10], followed by proteins that provide desiccation tolerance [11]. Key in desiccation tolerance are late embryogenesis (LEA) proteins including dehydrins, which accumulate in mature seeds before desiccation and are a hallmark of seed maturation [12–16]. LEA proteins are involved in controlling water loss, protecting cells during drying, and stabilising proteins in low water conditions [17,18]. Dehydrins are a specific class of LEA proteins that are involved in stress tolerance, including electrolyte leakage across membranes, lipid peroxidation, membrane binding and stabilisation, and cryoprotection of enzymes [19,20].

Late in maturation the seed is desiccated to 10-15% water, triggering dormancy [21]. Dormancy is an important trait because it provides mature plants resistance to pre-harvest sprouting, a condition in which seeds germinate in the field while still on the mother plant if exposed to moist conditions [16,22,23]. However, when dormancy is too strong, additional storage time is required prior to the malting process to pre-condition the seeds to a germinable state in order to attain the uniform germination desired [22,23]. In barley, this process is highly controlled in order to attain uniformly germinated grains with a high germination percentage [23–25].

The malting process involves three main steps: 1) Steeping, exposing the grain to periods of water soaking and aeration in order to raise the grain moisture content to about 35-45 %; 2) germination, carried out over 4 to 6 days with controlled relative humidity of 100 % and temperature (12 to 25 °C depending on grain type) in order to allow for the seedling growth and development; and 3) kilning, involving the drying of the seeds in large kilns in order to reduce the moisture content to about 4 to 6 % depending on the malt’s final use [2]. Kilning also stops enzymatic activity, preventing enzymes from fully breaking down proteins or starch, and stopping the embryo from using all of the released nutrients [3,26]. However, the malting process only works efficiently when all seeds germinate uniformly. Dormancy is therefore a severe problem in industrial barley malting [22–24].

Proteomics has been used previously in studies addressing many aspects of barley biology, such as identifying the proteins contained in a tissue or at a particular stage of development [26–32]. Recent years have seen increased use of proteomics to study the changes occurring in barley seeds during seed filling, grain maturation, and germination [3,33]. Mahalingam (2017), described the most complete proteomic analysis of mature barley seeds to date, identifying 1168 unique proteins. Comparatively, only 259 have previously been identified from mature barley seeds using 2-DE techniques. Out of these 1168 proteins identified, only 241 had meaningful UniProt annotations available [3], highlighting the need for additional functional characterisation of the barley proteome. A recent shotgun proteomics study monitored changes in the barley proteome during the malting process, identifying many large changes in the proteome [33].

Here, we used Sequential Window Acquisition of all THeoretical ions Mass Spectrometry (SWATH-MS) proteomic analysis of mature, germinated, and dormant barley seeds in a controlled laboratory germination environment to better understand the molecular features that drive the shift from mature barley seeds into germination in the presence of water. We also investigated the proteomic features of seeds that failed to germinate and their association with dormancy. Finally, we explored the proteomic profile of barley germination during industrial malting to verify the consistency of the observed proteomic changes across these massive differences in scale.

## Materials and Methods

### Laboratory Germination

Seeds used in this study were all from the one barley variety, Stirling. Stirling is an established malting grade barley variety with moderate to strong dormancy. Prior to imbibition, five seeds were selected as pre-imbibition controls. Five replicates of 100 seeds were then surface sterilized with 1% hypochlorite. These were rinsed thoroughly, air-dried, and placed in five 9 mm petri dishes lined with two Whatman #1 filter papers. Double distilled water (4 mL) was added and the dishes were covered and incubated in the dark at 22 °C. Seeds for proteomic analysis were selected on a single seed basis, where one seed was selected at random for each treatment category from each replicate dish, making five biological replicates for each treatment (Table 1). After sampling, each seed was placed in a separate protein LoBind tube (Eppendorf) and stored at −20 °C until extraction.

**Table 1.**
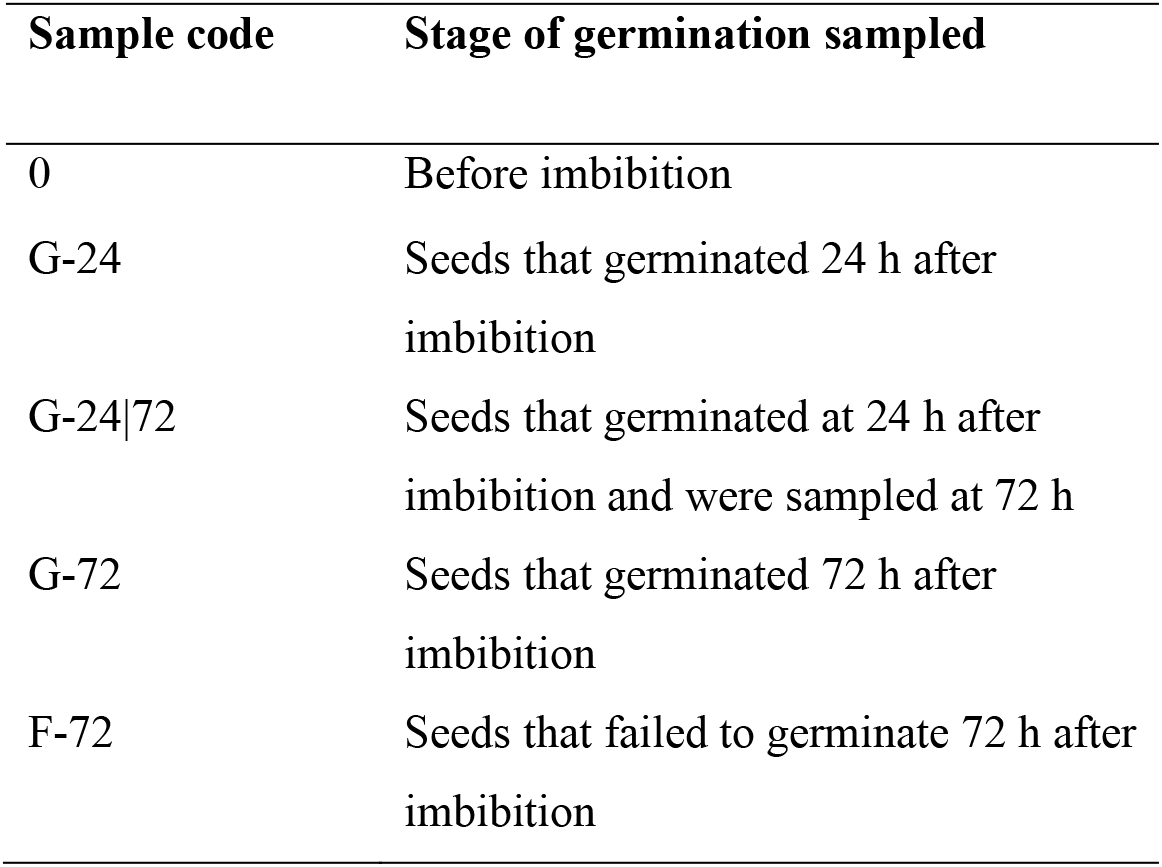
Seed sampling schedule.

### X-Ray visualisation

X-ray visualisation of randomly selected seeds was performed using a Faxitron X-Ray MX-20. Seeds were selected for visualization before imbibition (0), germinated at 24 h (G-24), 48 h post germination (G-24|72), and that failed to germinate 72 h after imbibition (F-72).

### Industrial Malting

Malting samples were collected from two commercial malt facilities as previously described [34], consisting of a mix of Buloke and Gairdner barley varieties. Both facilities performed a malting program of 24 h steeping, 88 h germination, and kilning to 80 °C. Malt samples were collected, frozen, and air-dried until under 10% moisture if wet. Rootlets and acrospires were removed before storage. Samples from both malt facility 1 (Malt 1) and malt facility 2 (Malt 2) were mature/pre-imbibition seeds (0), 24 h after imbibition (G-24), 48 h after imbibition (G-48), and 72 h after imbibition (G-72). Six biological replicates were included for each treatment.

### Protein extraction

Proteins from barley seeds were extracted and prepared for mass spectrometry as previously described [31]. Individual seeds were ground using a mortar and pestle to yield a homogenous powder, then placed in 1.5 mL Protein Lobind tubes (Millipore). To each sample, 600 μL of 50 mM Tris-HCl buffer pH 8, 6 M guanidine hydrochloride, and 10 mM DTT was added, mixed thoroughly by vortexing and incubated at 37 °C for 30 min with shaking. Cysteines were alkylated by addition of acrylamide to a final concentration of 30 mM and incubation at 37 °C for 1 h with shaking. Excess acrylamide was quenched by addition of DTT to an additional final concentration of 10 mM. Samples were centrifuged at 18,000 rcf for 10 min and 10 μL of the supernatants precipitated by addition of 100 μL 1:1 methanol:acetone and incubation at - 20 °C overnight. Precipitated proteins were pelleted by centrifugation at 18,000 rcf for 10 min and 1 min, with the supernatant being discarded after each centrifugation. The protein pellets were then air dried, resuspended in 100 μL of 100 mM ammonium acetate with 0.5 μg trypsin (Proteomics grade, Sigma) and incubated at 37 °C for 16 h with shaking.

### Mass spectrometry

Peptides were desalted with C18 ZipTips (Millipore) and measured by LC-ESI-MS/MS using a Prominence nanoLC system (Shimadzu) and TripleTof 5600 instrument with a Nanospray III interface (SCIEX) as previously described [35]. Approximately 1 μg or 0.2 μg desalted peptides, as estimated by ZipTip binding capacity, were injected for data dependent acquisition (DDA) or data independent acquisition (DIA), respectively. LC parameters were identical for DDA and DIA, and LC-MS/MS was performed essentially as previously described [36]. Peptides were separated with buffer A (1% acetonitrile and 0.1% formic acid) and buffer B (80% acetonitrile with 0.1% formic acid) with a gradient of 10–60% buffer B over 14 min, for a total run time of 24 min per sample. Gas and voltage setting were adjusted as required. For DDA analyses, an MS TOF scan from m/z of 350–1800 was performed for 0.5 s followed by DDA of MS/MS in high sensitivity mode with automated CE selection of the top 20 peptides from an *m/z* of 40–1800 for 0.05 s per spectrum and dynamic exclusion of peptides for 5 s after 2 selections. Identical LC conditions were used for DIA SWATH, with an MS-TOF scan from an *m/z* of 350–1800 for 0.05 s followed by high-sensitivity DIA of MS/MS from *m/z* of 50-1800 with 26 *m/z* isolation windows with 1 m/z window overlap each for 0.1 s across an m/z range of 400–1250. Collision energy was automatically assigned by the Analyst software (SCIEX) based on the centre of each *m/z* window range. Sample injection order was randomized.

### Data Analysis

Peptides and proteins were identified using ProteinPilot 5.1 (SCIEX), searching against High confidence proteins from transcripts from *Hordeum vulgare* IBSC v1 (downloaded 3 May 2017; Plant Genome and Systems Biology (PGSB), ftp://ftpmips.helmholtz-muenchen.de/plants/barley/genome_release2017; 248180 total entries), with settings: sample type, identification; cysteine alkylation, acrylamide; instrument, TripleTof 5600; species, none; ID focus, biological modifications; enzyme, trypsin; search effort, thorough ID. The proteins identified at 1% FDR by ProteinPilot were used to generate ion libraries for analysis of SWATH data. The abundance of peptides and proteins were determined using PeakView 2.1 (SCIEX) as previously described [31,35,37]. PeakView 2.1 (SCIEX) was used for analysis with the following settings: shared peptides, allowed; peptide confidence threshold, 99%; peptides per protein, 6; transitions per peptide, 6; false discovery rate, 1%; XIC extraction window, 6 min; XIC width, 75 ppm. The mass spectrometry proteomic datasets are available through the ProteomeXchange Consortium via the PRIDE partner repository with the dataset identifiers PXD019383 and PXD019384 for laboratory germination and industrial malting, respectively [38]. Proteins identified were matched against UniProtKB (downloaded 2 December 2017; 555318 total entries), using BLAST+ to find the best matching annotated entry. For protein-centric analyses, protein abundances were re-calculated by removing all peptide intensities that did not pass a local FDR cut-off of 1%, processing each sample independently, using a python script (Supplementary Material – Recalculate MS). Protein abundances were recalculated as the sum of all peptides from that protein, and normalised to the total protein abundance in each sample. PeakView output was reformatted with a python script (https://github.com/bschulzlab/reformatMS), applying a peptide FDR cut-off of 1% to remove ion measurements for low quality peptides from each sample, and reformatting appropriate for use with MSstats [31]. Differences in protein abundance were compared with a linear mixed model using MSstats (2.4) in R [39], with Benjamini-Hochberg corrections adjusting for multiple comparisons and a significance threshold of P = 10^−5^, as previously described [31]. Gene ontology (GO) term enrichment was performed using GOstats (2.39.1) in R [40] applying Bonferroni correction and a significance threshold of P = 0.05. Proteins and samples were both clustered with Cluster 3.0, applying a hierarchical, uncentered correlation, and complete linkage [41].

## Results

We aimed to investigate how germination affects the barley seed proteome and gain molecular insights into the causes of dormancy. X-ray imaging of selected seeds provided physical validation that germination had proceeded as expected in each of the groups of seeds (Fig. 1B). Before imbibition (0), when seeds had not commenced germination, they appeared intact with a darker endosperm than the other seed parts (Fig. 1B). Seeds that had germinated at both 24 h and 72 h (G-24 and G-72, respectively) appeared swollen with clear visible signs of germination; the radicle could be seen emerging from the seed (Fig. 1B). For seeds that germinated at 24 h and were sampled at 72 h (G-24|72), the roots were elongated and the plumule was emerging (Fig. 1B). The seeds which failed to germinate by 72 h (F-72) showed no visible signs of germination apart from appearing swollen (Fig. 1B). By 24 h post imbibition 82% of seeds had germinated, by 48 h post imbibition 93% of seeds had germinated, and by 72 h post imbibition 97% of seeds had germinated.

**Figure 1.**
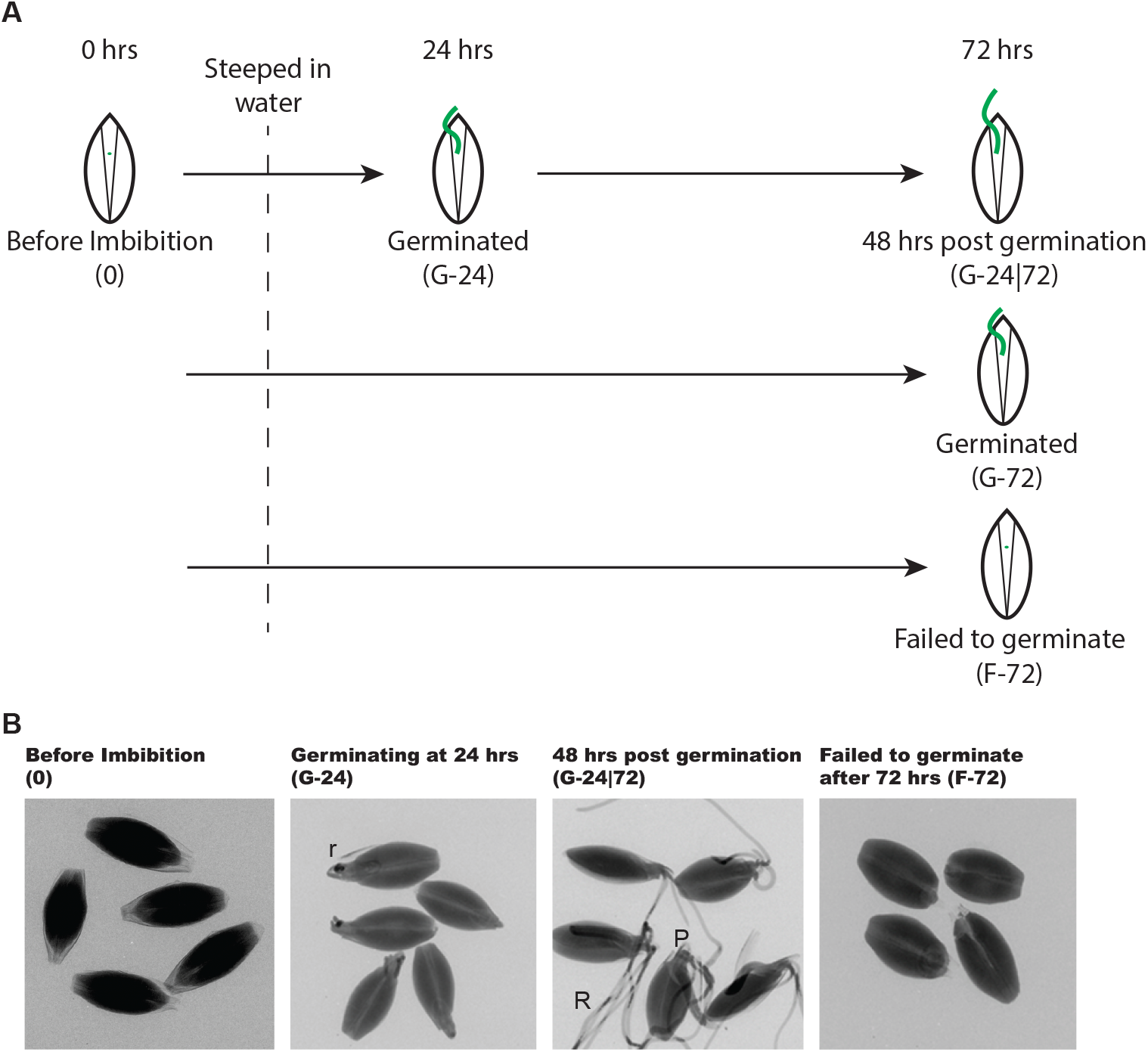
Overview of experimental design for laboratory germination. **(A)** Seed samples were taken at each point shown for DIA/SWATH-MS proteomics. **(B)** X-ray images of barley seeds at each sampled time point. r, radicle; R, roots; P, plumule.

To study the molecular details of the germination process, we performed DIA/SWATH-MS proteomics on barley seeds before imbibition (0), seeds germinating at 24 h (G-24), seeds germinating at 72 h (G-72), seeds that germinated at 24 h and were sampled at 72 h G-24|72), and seeds that failed to germinate by 72 h (F-72) (Fig. 1). A total of 209 proteins were identified and quantified (Table S1), and the results revealed major changes in the barley seed proteome during germination and also in the seeds which failed to germinate.

### Variation and differences present in the barley seed proteome

To investigate how barley seeds changed after imbibition, throughout germination, and when seeds are unable to germinate, we first examined the overall variance in protein abundance between samples. When comparing the proteomes of seeds at different stages of development we found that while their proteomes were generally quite similar, there were noticeable, subtle differences (Fig. 2A). Clustering of the seed proteomes showed that samples clustered by stage of development (Fig. 2A). Principal component analysis (PCA) provided an overview of the proteomic variability between samples (Fig. 2B). The PCA showed partial separation and clustering of before imbibition samples (blue) from other groups. We also observed partial separation of failed to germinate samples (grey) from samples that did germinate (Fig. 2B). To gain insight into the differences in the seed proteomes at various times post-imbibition, we statistically compared each condition to seeds before imbibition (0) (Fig. 2C and Table S2). A large number of proteins showed significant differences in abundance when compared to seeds before imbibition (0), the majority of which were significantly different for all stages of development (Fig. 2C and Table S2).

**Figure 2.**
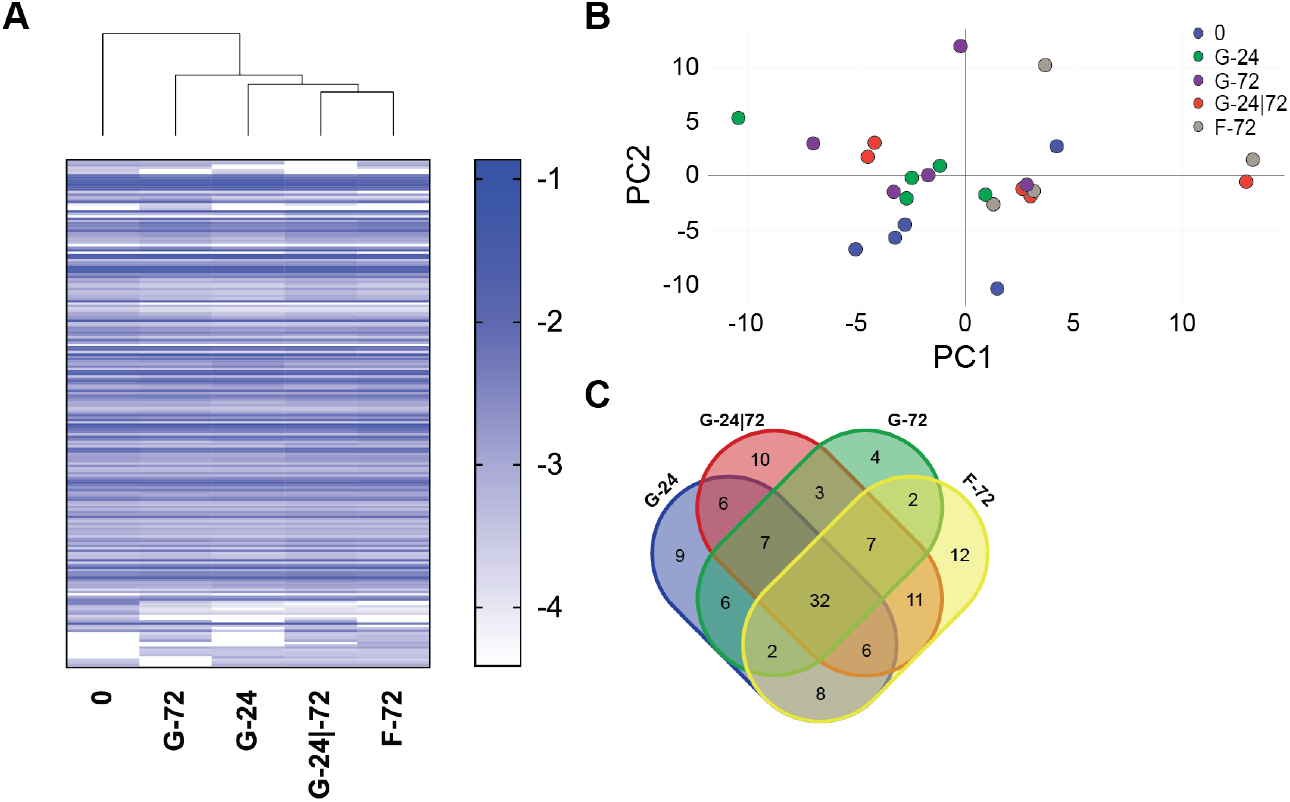
The effect of germination, growth, or lack of germination on the barley seed proteome. **(A)** Heat map of log_10_ protein abundance normalised to total protein abundance in each sample. Samples were clustered using Cluster 3.0, shown as a dendrogram; each horizontal line represents one of the total 209 proteins measured. **(B)** Principal component analysis (PCA) of the log_10_ protein abundances normalised to total protein abundance in each sample. Coloured by development stage: before imbibition (O), blue; germinated at 24 h (G-24), green; germinated at 72 h (G-72), purple; 48 h post germination (G-24|72), red; failed to germinate 72 h after imbibition (F-72), grey. The first component (x-axis) accounted for 13.57% of the total variance and the second (y-axis) 10.75%. **(C)** Venn Diagram of the number of proteins significantly (P<10^−5^) different in abundance compared to seeds prior to imbibition (0). Values, mean of biological replicates n=5. Error bars, SEM.

### Imbibition triggers a shift in the proteome away from stress response and lipid storage

Germination of barley seeds involves many complex processes, including development of the embryo, emergence of the radicle from the seed, degradation of the endosperm cell wall, and digestion of seed storage proteins and starch. These processes of germination are therefore likely to reflect a rich and dynamic proteome. To gain insights into these changes, we compared the proteomes of pre-steeped seeds (0), seeds germinated at 24 h (G-24), and seeds 48 h post germination (G-24|72) (Table S3). This analysis revealed that germination strongly altered the proteome of barley seeds (Fig. 3A). A suite of late embryogenesis proteins (LEA) proteins were found to be the most differentially abundant proteins between germinating seeds and pre-steeped seeds (Fig. 3B). Four proteins belonging to the LEA family were significantly less abundant in all germinating seeds compared to pre-steeped seeds (Fig. 3B). We also identified two dehydrins, DHN3 and DHN4, that significantly decreased in abundance after imbibition (Fig. 3B).

**Figure 3.**
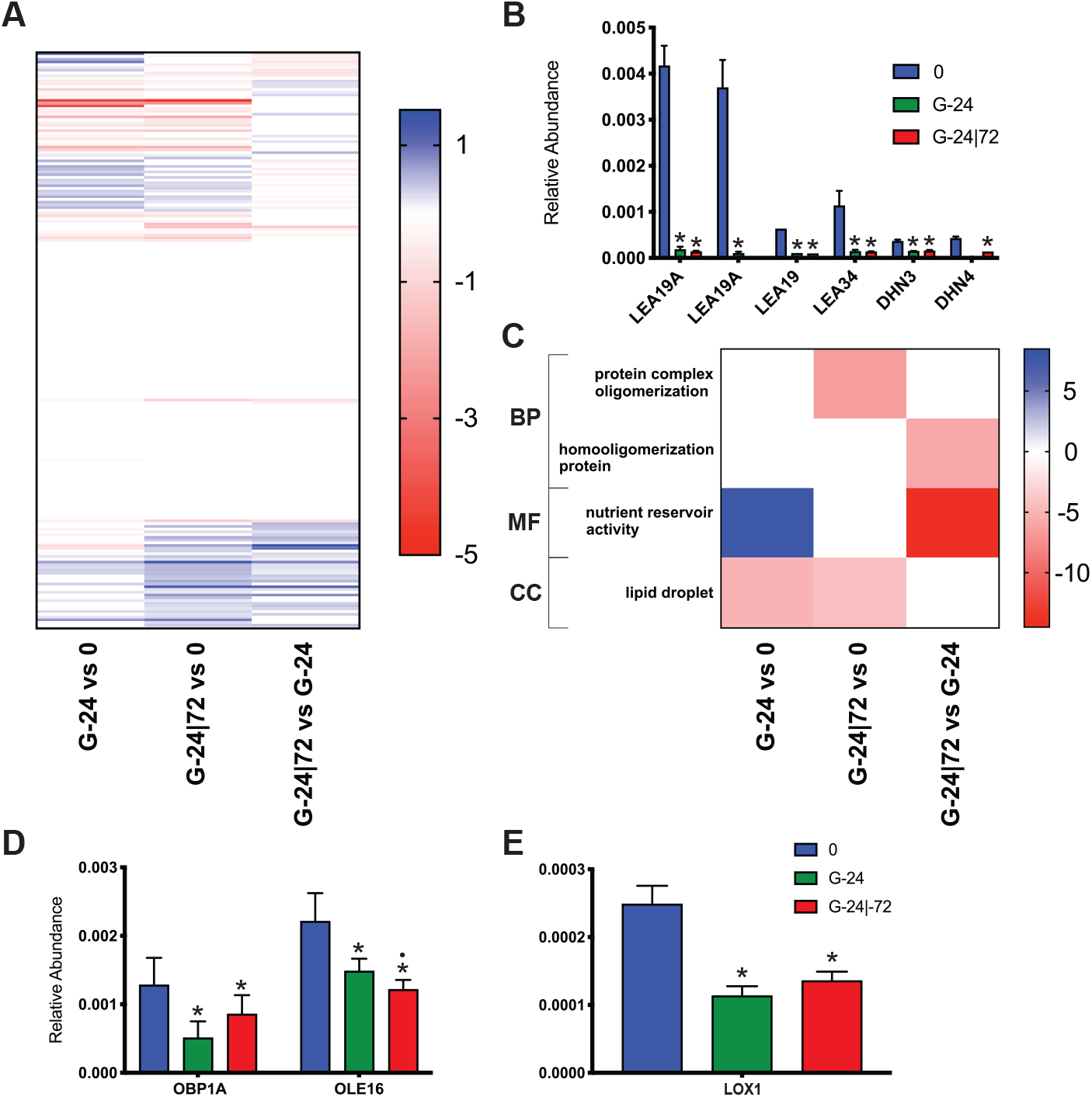
The effect of germination on the barley seed proteome. **(A)** Heat map of significantly differentially abundant proteins. Values shown as log2 (fold change) for proteins with significant differences in abundance between G-24 to 0, G-24|72 to 0, or G-24|72 to G-24 (P<10^−5^). **(B)** Normalised relative abundance of late embryogenesis proteins; two forms of LEA19A, LEA19, LEA34, DHN3, and DHN4. **(C)** Heat map of significantly enriched GO terms identified by GOstats analysis. Values shown as −log_2_ of Bonferroni corrected p-value for GO terms which were significantly enriched (p < 0.05) in each comparison. Values are shown as positive values for GO terms enriched in proteins more abundant in 0, 0, or G-24, in each respective comparison; values are shown as negative values for GO terms enriched in proteins more abundant in G-24, G-24|72, or G-24|72, in each respective comparison. **(D)** Normalised relative abundance of two lipid associated proteins, OBP1A and OLE16. **(E)** Normalised relative abundance of LOX1. Values, mean of biological replicates n=5. Error bars, SEM. * p < 10^−5^ with comparison to 0. •p < 10^−5^ with comparison between G-24 and G-24|72.

Following statistical comparison of protein abundance, we performed GO term enrichment analysis associated with these changes to gain an understanding of the key processes that changed in the germinating seeds (G-24 and G-24|72) compared to pre-steeped seeds (0) (Fig. 3C and Table S4). “Lipid droplet” was significantly enriched in proteins significantly less abundant in G-24 and G-24|72 compared to 0 (Fig. 3C). Two proteins contributed to the GO term enrichment of “Lipid droplet”, Oil body-associated protein 1A (OBP1A) and Oleosin 16 kDa (OLE16), which are involved in regulating oil body size and localisation and stabilizing lipid bodies during desiccation, respectively [42,43]. (Fig. 3D). Both OBPIA and OLE16 were found to be significantly more abundant before imbibition in 0 compared to germinated seeds at both G-24 and G-24|72 (Fig. 3D). Alongside the GO term enrichment of “Lipid droplet”, we found a significantly higher abundance of Linoleate 9S-lipoxygenase 1 (LOX1) in mature seeds (0) compared to germinating seeds (G-24 and G-24|72) (Fig. 3E). LOX1 is involved in oxylipin biosynthesis, a part of lipid metabolism. “Protein complex oligomerization” was significantly enriched in 0 compared to G-24|72 and “homooligomerization protein” was significantly enriched in G-24 compared to G-24|72 (Fig. 3C). Cupincin (CUCIN) and 16.9 kDa class I heat shock protein 2 (HS16B) were associated with both “protein complex oligomerization” and “homooligomerization protein” terms, while 23.2 kDa heat shock protein (HS232) and 17.8 kDa class II heat shock protein (HSP22) were only associated with “protein complex oligomerization” (Fig. S1). Cupincin, belonging to the cupin superfamily, is a probable zinc metallo-protease [44]. HS16B, HS232, and HSP22 are all involved in the heat shock response system as protein folding or stabilisation chaperones. Finally, “nutrient reservoir activity” was significantly enriched in G-24 compared to 0 and G-24|72 (Fig. 3C). A suite of seed storage and related proteins were associated with the “nutrient reservoir activity” GO term: AVLA1, AVLA4, CUCIN, GLT3, HOG1, HOG3, HOR1, HOR7, SSG1, HOR3, SPZ4, and VSPA (Fig. S1). This enrichment pattern describes an initial increase in the abundance of enzymes and proteins associated with “nutrient reservoir activity” after imbibition as germination begins, followed by a decrease in the abundance of nutrient reservoirs as seed development continues (Fig. 3C and S1), consistent with induction and activation of systems for nutrient mobilisation.

### Seed dormancy leads to enrichment of chromatin related processes

In an industrial malting setting, the failure of seeds to germinate under standard conditions requires additional storage to pre-condition the seeds into a germinable state. This additional storage is needed to attain uniform germination in the malting process. To obtain insights into the mechanisms underlying dormancy, we compared the proteome of seeds that failed to germinate (F-72) to pre-steeped (0) and germinated seeds 72 h post imbibition (G-72 and G-24|72) (Table S5). This analysis showed that, as expected, a large number of proteins were significantly different in seeds that failed to germinate (F-72) compared to germinating seeds (G-72 and G-24|72) (69 and 49 proteins) (Fig. 4A). Surprisingly, however, we also detected large changes in seeds that failed to germinate (F-72) compared to mature seeds (0) (80 proteins) (Fig. 4A), indicating that even though the seeds that failed to germinate did not show gross signs of germination (Fig. 1B), they did have substantial biochemical activity.

**Figure 4.**
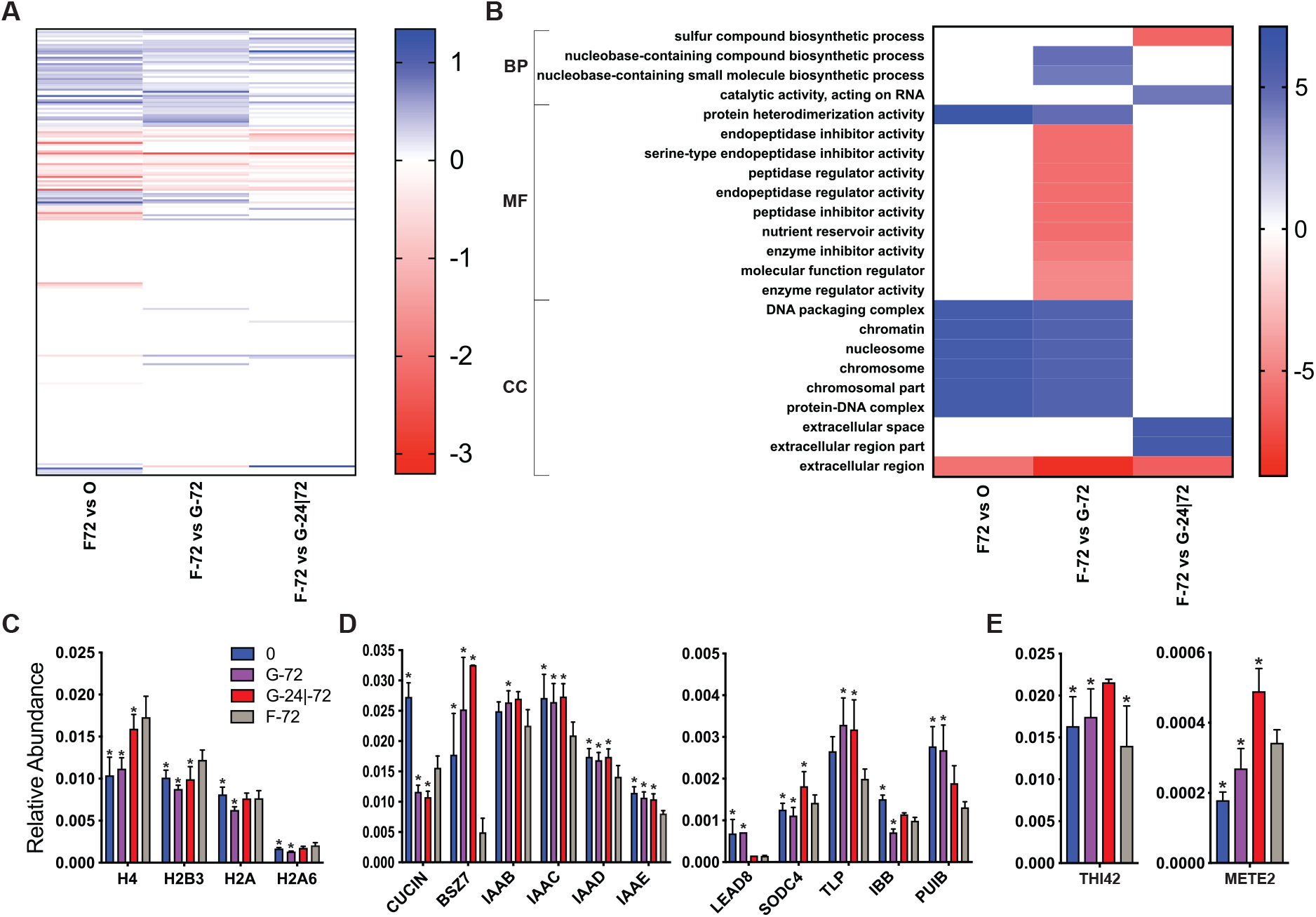
Failure to germinate has significant effects on the barley seed proteome. **(A)** Heat map of significantly differentially abundant proteins. Values shown as log2 (fold change) for proteins with significant differences in abundance between F-72 to 0, F-72 to G-72, and F-72 to G-24|72 (P<10^−5^). **(B)** Heat map of significantly enriched GO terms identified by GOstats analysis in each comparison. Values shown as −log_2_ of Bonferroni corrected p-value for GO terms which were significantly enriched (p < 0.05). Values. Are shown as positive values for GO terms enriched in proteins more abundant in F-72; values are shown as negative values for GO terms enriched in proteins less abundant in F-72. Biological process, BP; molecular function, MF; chemical compartment, CC. **(C)** Abundance of H4, H2B3, H2A, and H2A6, proteins involved in histone formation. **(D)** Abundance of CUCIN, BSZ7, IAAB, IAAC, IAAD, IAAE, LEAD8, SODC4, TLP, IBB, and PUIB. **(E)** Abundance of THI42 and METE2, proteins involved in sulfur compound biosynthesis process. Values, mean of biological replicates n=5. Error bars, SEM. * p < 10^−5^ with comparison to 0. • p < 10^−5^ with comparison against F-72.

We performed GO term enrichment analysis to gain an understanding of underlying processes associated with seeds that failed to germinate (Fig. 4B) (Table S6). Proteins that were more abundant in F-72 seeds that failed to germinate, compared to both 0 and G-72, were associated with GO terms involving DNA complexes, chromosome components, nucleotide biosynthesis and protein heterodimerization (Fig. 4B). These GO terms were associated with the higher abundance of histones (H2A6, H2A, 2B3, H4) in F-72 compared to 0 and G-72 (Fig. 4B, 4C, and Table S6). This higher apparent abundance of histones could be due to an actual increase in histone protein abundance in F-72 seeds that failed to germinate. Alternatively, or in addition, changes in the post-translational landscape of histones might also affect the abundance of tryptic peptides. This is particularly relevant given regulated changes in the pattern of histone acetylation and methylation on lysine residues are critical in controlling germination [45].

Proteins that were less abundant in F-72 seeds that failed to germinate compared to G-72 were enriched in GO terms for “peptidase regulator/inhibitor activity”, and compared to 0, G-72, and G24|72 for “extracellular region” (Fig. 4B and Table S6). The proteins that contributed to all of these GO terms were predominantly alpha-amylase/trypsin inhibitor (chloroform/methanol-soluble) proteins (Fig. 4D and Table S6). This class of protein were universally less abundant in F-72 relative to all other stages of germination, including 0 seeds before imbibition (Fig. 4D).

A particularly interesting enriched biological process was “Sulfur compound biosynthetic process”, enriched in proteins that were significantly less abundant in F-72 compared to G-24|72 (Fig. 4B). This enrichment was driven by the significantly higher abundance of Thiamine thiazole synthase 2 (THI42) and 5-methyltetrahydropteroyltriglutamate-homocysteine methyltransferase 2 (METE2) in germinating seeds (G-24|72) compared to those that failed to germinate (F-72) (Fig. 4E).

In summary, F-72 seeds that failed to germinate had a proteome profile that demonstrated the presence of substantial biochemical activity distinct from that of seeds undergoing germination. This was consistent with results from PCA (Fig. 2B), which showed partial clustering of F-72 seeds distinct from both germinating seeds and seeds before imbibition.

### Industrial scale malting shows similar proteome changes to laboratory germination

Malting of barley seeds for brewing is a three-step process: steeping, germination and kilning. To determine if the proteomic changes we detected in laboratory-scale germination were also relevant at industrial scale, we performed DIA/SWATH-MS proteomics on samples taken throughout the malting processes at two independent commercial malt houses (Malt 1 and Malt 2) [34]. We studied barley seeds from both malt houses collected from before imbibition (0), 24 h after imbibition (G-24), 48 h after imbibition (G-48), and 72 h after imbibition (G-72). We first compared the overall proteome of seeds germinating at the two malt houses. This analysis showed that the two malt houses had similar profiles; at each step throughout the malting process the proteomes of barley seeds at the two malt houses were very similar (Fig. 5A and 5B). Unguided clustering of the germinating malt proteomes showed that samples generally clustered by stage of malting rather than by facility (Fig. 5A). Samples from early stages of malting, G-24 and G-48, clustered by malthouses rather than stage of malting (Fig. 5A), however all four of these proteomes were very similar (Fig. 5B). PCA also showed that samples clustered strongly by the stage of malting (Fig. 5B).

**Figure 5.**
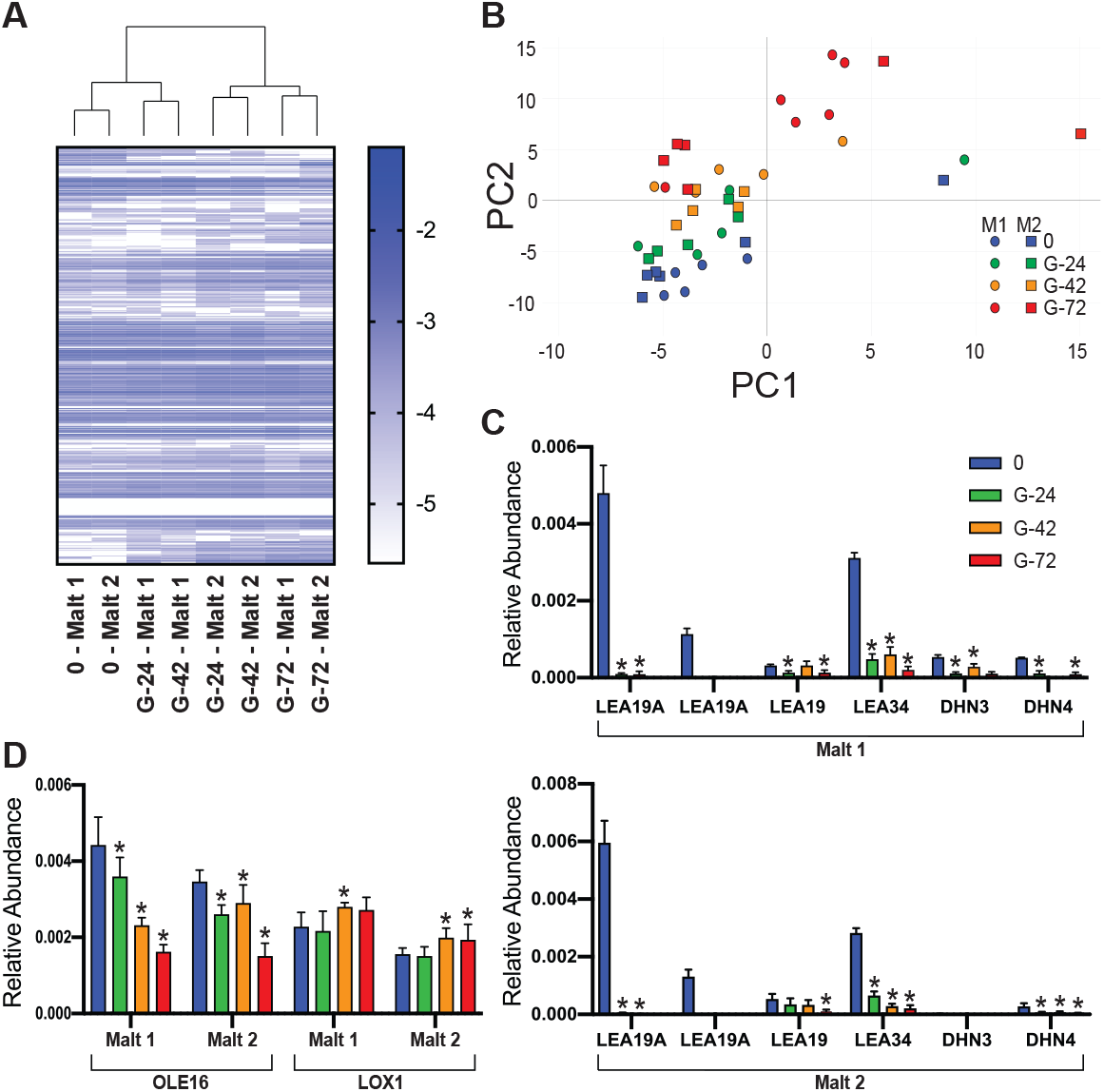
The proteomes of industrial malt changes as germination progresses. **(A)** Heat map of log_10_ of proteins normalised to total protein abundance and averaged across replicates. Samples were clustered, shown as a dendrogram. **(B)** Principal component analysis (PCA) of the −log_10_ of proteins values normalised to total protein abundance in each sample. The first component (x-axis) accounted for 20.52% of the total variance and the second (y-axis) 7.05%. **(C)** Abundance of late embryogenesis proteins, two LEA19As, LEA19, LEA34, DHN3, and DHN4 in both Malt 1 (top) and Malt 2 (bottom). **(D)** Abundance of OLE16 and LOX1. Malt 1, M1; Malt 2, M2; prior to imbibition, 0; 24 h after imbibition, G-24; 48 h after imbibition, G-48; 72 h after imbibition, G-72. Values, mean of biological replicates n=6 Error bars, SEM. * p < 10^−5^ with comparison to 0.

We next focussed on the changes in the proteome we had observed during germination at a laboratory scale (Fig. 3 and Table S7). LEA proteins were abundant in mature seeds, and showed a very large drop in abundance at all stages of germination post-imbibition at laboratory scale (Fig. 3B). We investigated the abundance of LEA proteins during malting at industrial scale, and found that at this scale they also showed significant reductions in abundance during malting at the two independent malt houses (Fig. 5C). Two forms of LEA19A, LEA19, LEA34, DHN3, and DHN4 were all significantly lower in abundance after imbibition and during malting in both facilities (Fig. 5C). Oleosin (OLE16) decreased in abundance during germination at laboratory scale, and this was also verified at industrial scale at both malt houses (Fig. 5D). Linoleate 9S-lipoxygenase 1 (LOX1) decreased in abundance during germination at laboratory scale, but was found to increase in abundance during malting at industrial scale at both malt houses (Fig. 5D).

## Discussion

Germination is integral to the development of mature barley seeds into malt for use in the beer brewing process. The changes that occur to the seed and its proteome during germination are important in allowing efficient mashing; ensuring the availability of sugars, amino acids, and other nutrients for yeast fermentation; and also affect the proteins, carbohydrates, and other molecules remaining in the final beer. The failure of seeds to predictably germinate in an industrial malting setting can cause substantial inefficiencies in the malting process and quality problems downstream in the brewing process. We observed dramatic changes in the proteome of barley seeds during laboratory scale germination (Fig. 2 and 3) which were consistent with industrial scale malting (Fig. 5). We also observed substantial changes in the proteome of seeds that failed to germinate, consistent with substantial but ultimately ineffective biochemical activity (Fig. 4 and 6).

**Figure 6.**
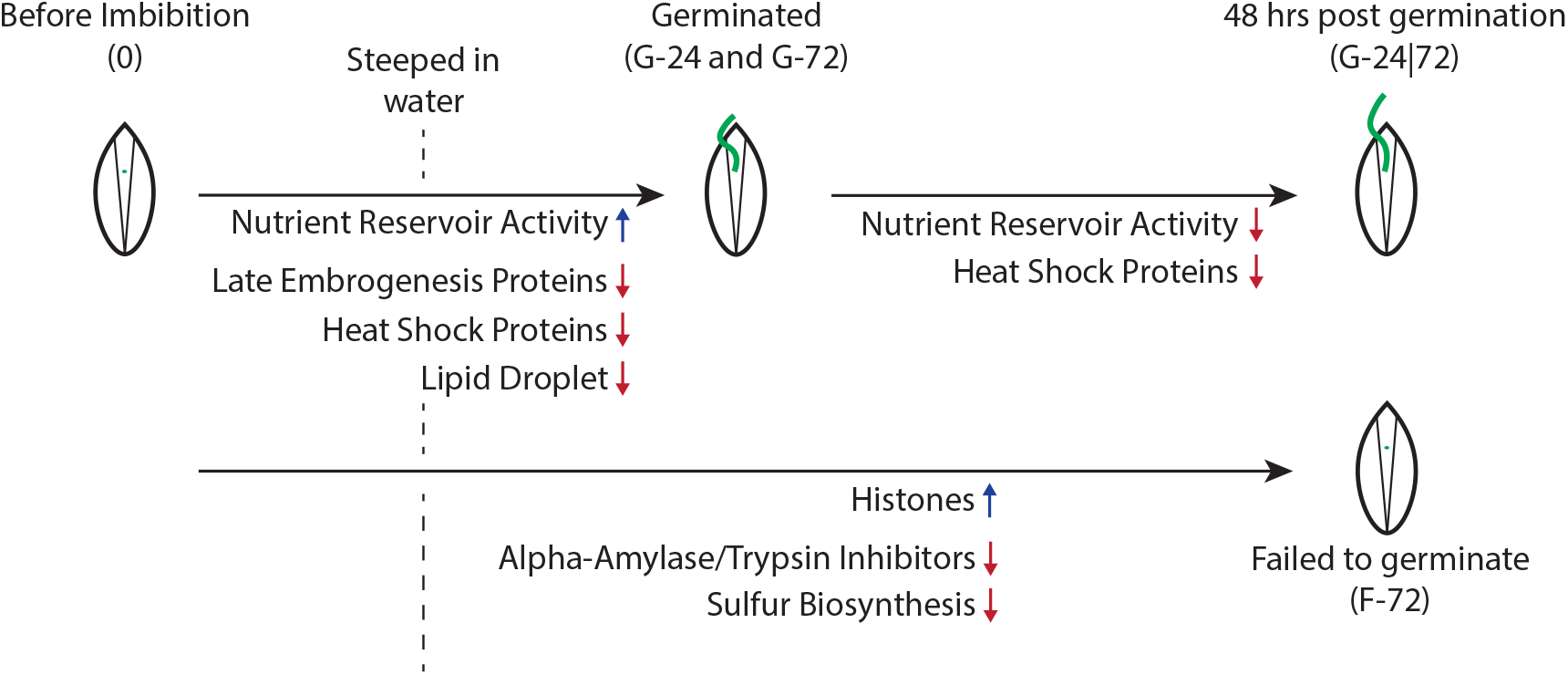
Overview of changes to the barley seed proteome during germination and dormancy.

Imbibition resulted in a rapid change to the barley proteome. Several groups of proteins were dramatically reduced in abundance within 24 h of imbibition, many of which were involved in protecting the desiccated barley seed [12–16,46,47]. In particular, the striking loss of late embryogenesis proteins (LEA) in germinating seeds compared to seeds before imbibition (Fig. 2B and 5C) is likely linked to the importance of these proteins in desiccation tolerance [12–16,46,47]. Dehydrins, a subgroup of LEA proteins [48,49], also showed a dramatic decrease in abundance post-imbibition (Fig. 2B and 5C). These LEA proteins are likely critical for desiccation tolerance during seed storage, but are no longer required once germination commences. The mechanism of LEA protein degradation is unclear, but likely requires specific and controlled proteolysis. Imbibition also resulted in a significant loss of “lipid droplet” associated proteins (Fig. 3D and Table S4), consistent with mobilisation of lipid stores for use by the developing embryo. Low levels of oil body-associated protein 1A (OBP1A) in mature maize seeds have been found to significantly decrease germination rate, highlighting its importance in germination [42]. Linoleate 9S-lipoxygenase 1 (LOX1) is involved in the biosynthesis of oxylipins, a group of oxygenated fatty acid derivatives which accumulate during maturation and inhibit germination [50]. Together, this reduction of proteins involved in lipid storage and oxylipin production after imbibition is consistent with the commencement of germination. Proteins associated with “nutrient reservoir activity” increased in abundance early in germination and then decreased in abundance, likely linked to the mobilisation and utilisation of amino acid storage proteins through germination (Fig. 3C and Table S4).

We found that a small but substantial proportion of barley seeds failed to germinate under laboratory conditions. This is an established behaviour of stored barley seeds known as dormancy. While physiologically important to control premature germination, inappropriate levels of dormancy cause substantial inefficiencies and economic costs to industrial malting. To study the molecular correlates of dormancy, we compared the proteomes of seeds that failed to germinate with seeds before imbibition and after germination. This analysis revealed that histone proteins were enriched in seeds that failed to germinate compared to both mature seeds and germinating seeds (Fig. 4B). Dehydration and rehydration during seed maturation and imbibition, respectively, is associated with oxidative stress which can lead to DNA damage, resulting in loss of seed viability and associated with dormancy [51–53]. The abundance of histones in seeds that failed to germinate may be reflective of DNA damage. Alternatively, the apparent changes in the abundance of histones may be associated with changes in the chromatin post-translational landscape in these seeds. Methylation, acetylation, and other modifications of histone lysines are critical regulators of gene expression during germination [45]. These modifications may be perturbed in seeds that begin but fail to complete the process of germination. In turn, altered lysine modification may change the apparent abundance of peptides resulting from trypsin cleavage, which specifically cuts at lysine and arginine residues.

We also observed low abundance of proteins associated with “peptidase inhibitor/regulator activity” in seeds that failed to germinate, mainly alpha-amylase/trypsin inhibitor (chloroform/methanol-soluble CM) proteins. These proteins were lower in abundance in seeds that failed to germinate than seeds both before and after germination, suggesting that their low abundance may actively contribute to their failure to germinate. While these proteins protect the seed from fungal and insect attack [54,55], they are also able to inhibit barley proteases [56–58], and have been proposed to regulate aspects of germination [56–58]. They have also been reported to directly bind to chromatin [59], with potential roles in gene regulation consistent with the changes we observed in histone protein abundance in the seeds that failed to germinate.

Proteins associated with “Sulfur compound biosynthetic process” were low in abundance in seeds that failed to germinate. Sulfur is a critical element for plant growth, needed for the synthesis of methionine and cysteine. Both amino acids are not only required for protein biosynthesis, but are also integral in ethylene synthesis, biotin synthesis, and lysine catabolism [53]. One of the proteins we found that is associated with “Sulfur compound biosynthetic process”, was thiamine thiazole synthase 2 (THI42) [60]. THI42 transfers one of its own sulfides to a thiazole intermediate, a precursor of thiamine (vitamin B1) [60]. Thiamine phosphate derivatives are critical for a variety of cellular processes including the catabolism of sugars and amino acids. In addition to THI42, 5-methyltetrahydropteroyltriglutamate-homocysteine methyltransferase 2 (METE2), a catalyst in the formation of methionine, was also associated with “Sulfur compound biosynthetic process”. The increase during germination in the synthesis of methionine, cysteine, and downstream compounds required for efficient germination highlights the lack of sulfur compound synthesis in seeds that failed to germinate.

Our experimental design did not allow us to unequivocally differentiate between these differences in protein abundance being a cause or consequence of seeds failing to germinate. It is likely that the differences in sulfur metabolism were a consequence of a general lack of growth. It is possible that the differences in apparent histone abundance reflect DNA damage or epigenetic marks that actively prevent germination. Finally, the low abundance of alpha-amylase/trypsin inhibitor proteins in seeds that failed to germinate may reflect a biochemical deficiency in these seeds that reduced their germination efficiency.

To verify our findings at laboratory germination scale, we investigated key aspects of the proteome of barley seeds during germination at an industrial scale in two independent malt houses. A major change we observed in the barley seed proteome during germination at laboratory scale was a rapid loss of late embryogenesis related proteins after imbibition (Fig. 3B). We found that this large and significant change was also robustly observed in seed germinating at industrial malt house scale (Fig. 5C). We could also verify the decrease in the abundance of Oleosin 16 kDa (OLE16) during germination, consistent with mobilisation of lipid reservoirs. However, linoleate 9S-lipoxygenase 1 (LOX1), decreased in abundance after imbibition at laboratory scale, but increased in abundance during germination at malt house scale (Fig. 5D, 3D, and 3E). It has been previously reported that lipoxygenase activity increases during germination, and also that commercial malts have particularly high activity [61]. The discrepancy we observe between laboratory and industrial scales may be due to differences in the behaviour of barley seed germination at these different scales, or differences in the barley varieties that were used at laboratory and industrial scales. We observed somewhat higher variability in our laboratory scale experiments than at industrial scale, perhaps because we performed single-seed analyses at laboratory scale. We also note that enzyme abundance may not directly correlate with activity, as the latter can be influenced by localisation, activation, or other regulatory mechanisms.

In summary, this industry scale malting data shows that the key changes to the proteome we observed during germination are consistent from laboratory to industrial scales, validating the analysis of changes in the proteome of laboratory-scale barley germination as a model system for industrial malt house germination.

## Supplementary Material

### Supplementary Figures

**Figure S1.**
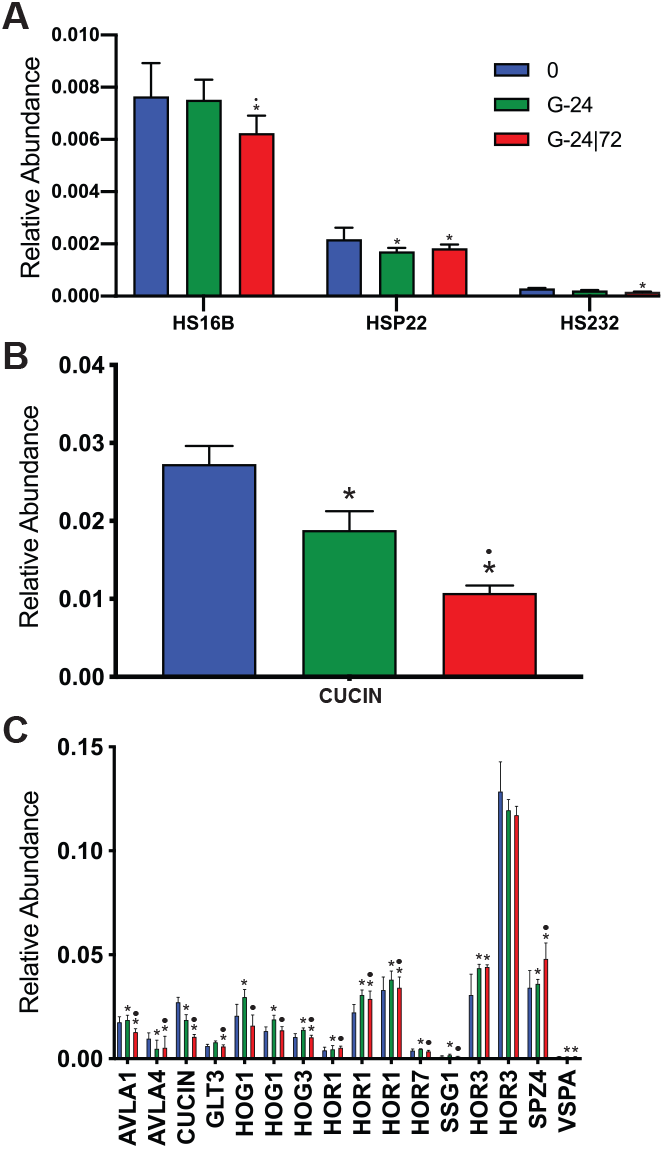
**(A)** Abundance of heat shock proteins; HS16B, HSP22, and HS232. **(B)** Abundance of CUCIN. **(E)** Abundance of seed storage and related proteins; AVLA1, AVLA4, CUCIN, GLT3, HOG1, HOG3, HOR1, HOR7, SSG1, HOR3, SPZ4, and VSPA. Values, mean. Error bars, SEM. * p < 10^−5^ with comparison to 0. • p < 10^−5^ with comparison between G-24 and G-24|72.

### Supplementary Table Legends

**Table S1. Protein SWATH Sheet**

**Table S2. MSstats vs 0**

**Table S3. Germination MSstats**

**Table S4. Germination GO term enrichment**

**Table S5. Failed to germinate MSstats**

**Table S6. Failed to germinate GO term enrichment**

**Table S7. Industry MSstats**

### Supplementary Material

#### Recalculate MS

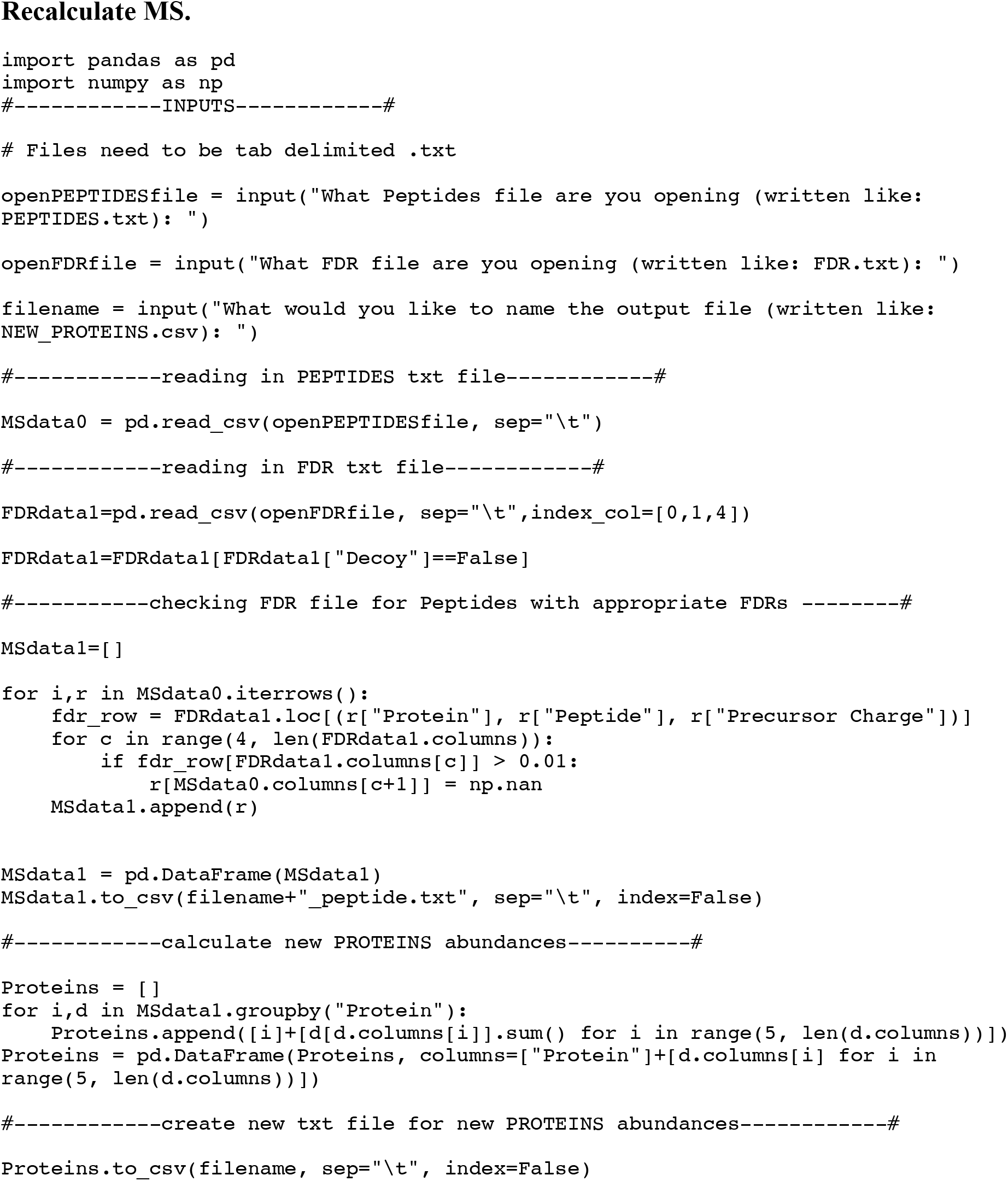

